## Supplementary Information for "Sensory sharpening and semantic prediction errors unify competing models of predictive processing in human speech comprehension"

### 1 Supplementary Figures

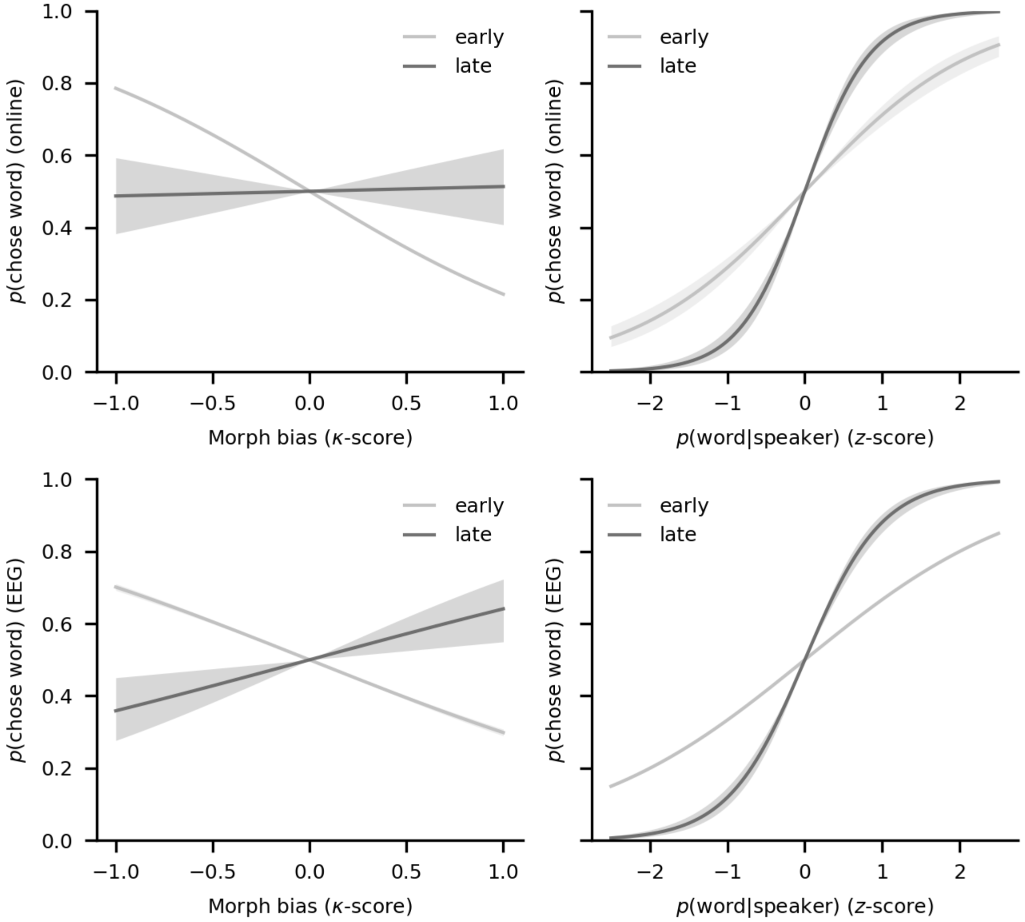


**Supplementary Fig. 1** Summary of key behavioural results from the online experiment one (top row) and EEG experiment two (bottom row). In both experiments, participants initially relied on remaining acoustic properties (i.e., morph bias to one of the words within each word pair), but decreased this reliance over time (left). Participants increasingly relied on the probability of the word given the speaker instead (right). Lines represent means, with shaded areas representing 95%-confidence intervals. For details, see Supplementary Table 1-2. Data and code supporting these findings are available from https://doi.org/10.17605/OSF.IO/SNXQM.

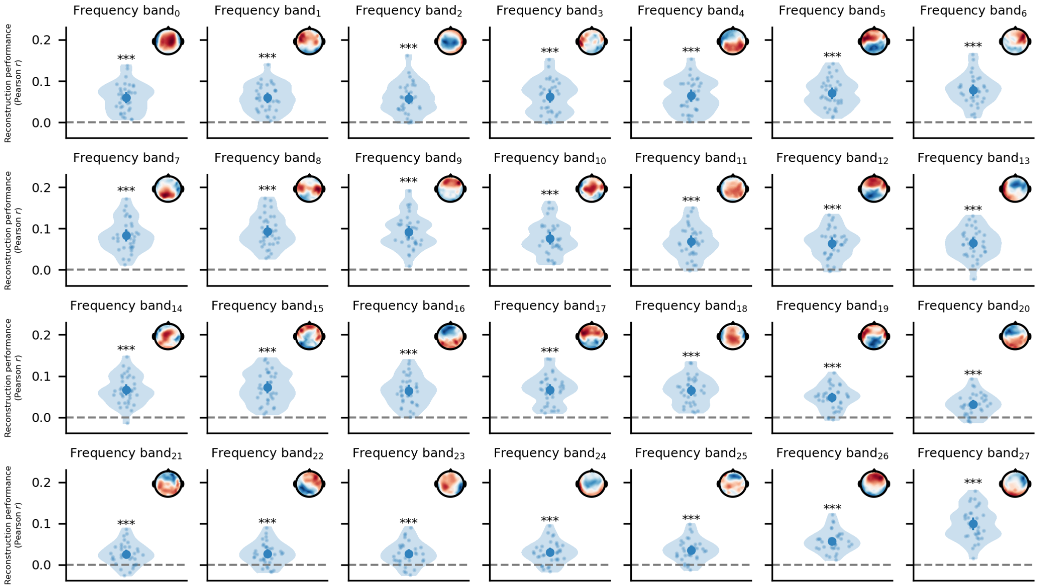


**Supplementary Fig. 2** Accuracy of stimulus reconstruction models[1] in all 28 individual frequency bands. Bold dots indicate means, with 95%-confidence intervals around them. Small dots represent individual participants. Inlaid topographies show the decoded pattern for this frequency band. ***, **, * represent $p \leq 1e^{-3}$, $p \leq 1e^{-2}$ and $p \leq 5e^{-2}$, respectively. For details, see Supplementary Table 3. Data and code supporting these findings are available from https://doi.org/10.17605/OSF.IO/SNXQM.

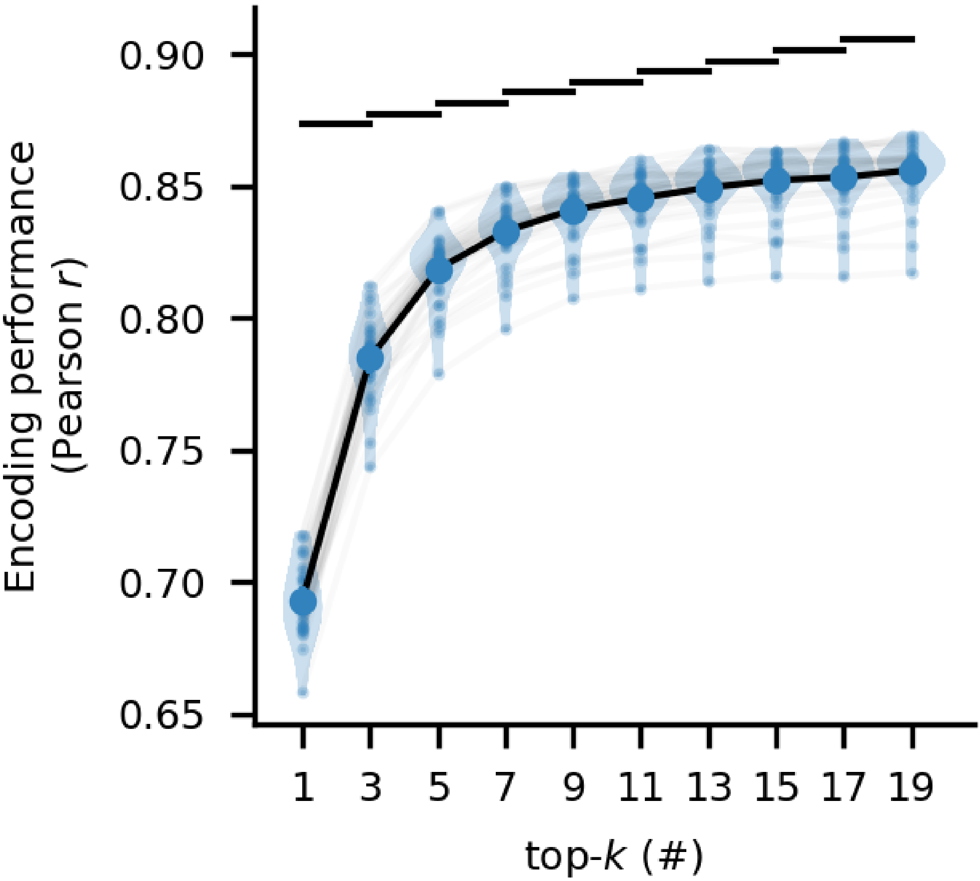


**Supplementary Fig. 3** We systematically refit models with increasing $k$ to test whether the brain makes a specific number of predictions $k$. Big dots represent group means, with 95%-confidence intervals around them. Small dots indicate individual participants. Big black bars on top indicate $p \leq 5e^{-2}$ for any contrast. For more details, see Supplementary Table 4. Data and code supporting these findings are available from https://doi.org/10.17605/OSF.IO/SNXQM.

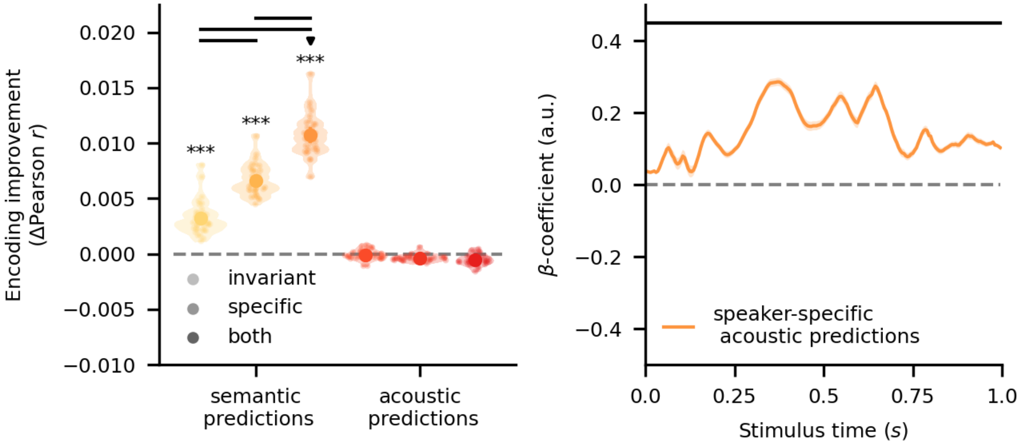


**Supplementary Fig. 4** Similarity encoding models using $k = 19$ predictions, because increasing the number of predictions $k$ yielded significant improvements in performance. Left: Results show that both speaker-invariant and speaker-specific acoustic predictions improve model performance, with the best model incorporating both at once. Critically, purely semantic predictions failed to improve performance. This is in line with our results based on $k = 5$, reported in Fig. 2. Big dots represent group means, with 95%-confidence intervals around them. Small dots represent individual participants. ***, **, * indicate $p \leq 1e^{-3}$, $p \leq 1e^{-2}$, and $p \leq 5e^{-2}$, respectively. The downward-facing triangle marks the best model overall. Big black bars between groups indicate $p \leq 5e^{-2}$. Right: Coefficients for speaker-specific acoustic predictions showed, again, that there was significant sharpening of neural representations, with a cluster between $0ms$-$1000ms$. Lines indicate mean, with shaded areas around them representing 95%-confidence intervals. Big black lines indicate $p \leq 5e^{-2}$. Data and code supporting these findings are available from https://doi.org/10.17605/OSF.IO/SNXQM.

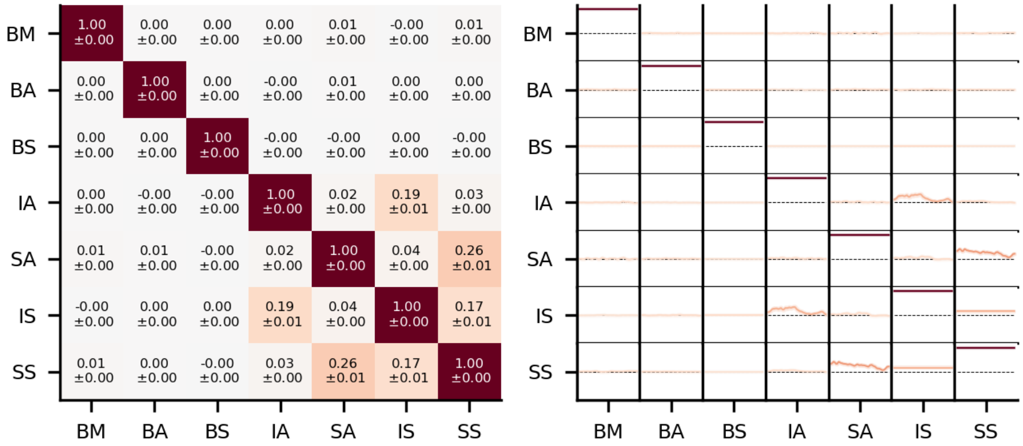


**Supplementary Fig. 5** Correlations between all hypothesis cRSMs used in regressing sensory cRSMs for top-$k$ predictions where $k = 5$. Hypothesis cRSMs included baseline morphs (BM), baseline acoustic predictions (BA), baseline semantic predictions (BS), speaker-invariant acoustic predictions (IA), speaker-specific acoustic predictions (SA), speaker-invariant semantic predictions (IS), and speaker-specific semantic predictions (SS). Baseline predictors were included in all models to control for acoustic properties of the morph (BM) as well as general acoustic (BA) and semantic predictions (BS) that were irrespective of our experiment. Left: Correlations were computed between all hypothesis cRSMs and averaged over time points. Individual cells contain means and standard errors computed over subjects. Right: Correlations were computed between all hypothesis cRSMs, but not averaged over time points. In each cell, time points (0.0-1.0$s$) are plotted on the x-axis, whereas correlations (-1 - 1) are plotted on the y-axis. Per cell, we plot chance level (dashed lines) and mean over subjects (solid lines) with confidence intervals indicated by shaded areas around means. Data and code supporting these findings are available from https://doi.org/10.17605/OSF.IO/SNXQM.

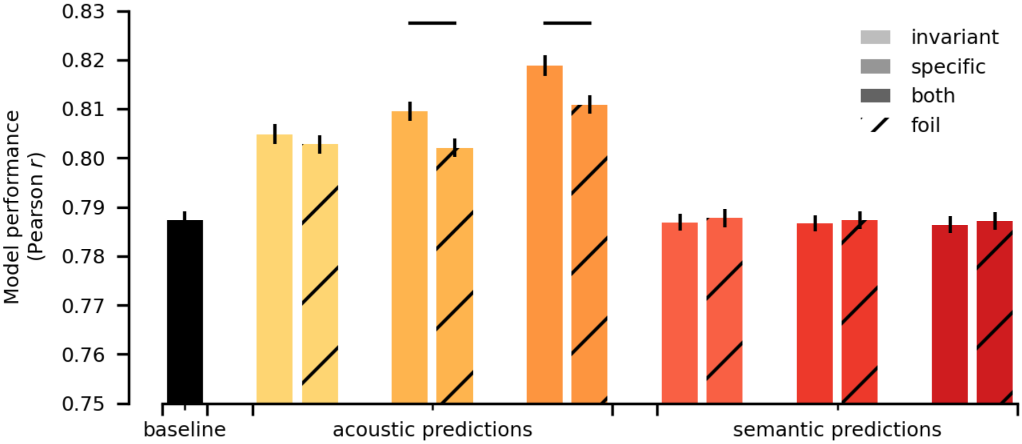


**Supplementary Fig. 6** Validation analysis using foil priors that disrupted participant-specificity of semantic priors. To confirm that performance of top-$k$ acoustic predictions depended on the trial-specific top-$k$ predictions, we compared performance of predictions from the true speaker-invariant and -specific semantic priors with coherent but not trial-specific foil semantic priors in cRSM regressions (see Validating top-$k$ acoustic predictions). In accordance with our main analysis, we set $k = 5$. As expected, this confirmed that speaker-specific acoustic predictions depended on the trial-specific predictions, indicating that they captured meaningful acoustic predictions. Note that, for visual clarity, we restricted the y-axis to a smaller range. Individual bars represent group means, with black lines indicating 95%-confidence intervals around the mean. Big bars between groups indicate $p \leq 5e^{-2}$. Data and code supporting these findings are available from https://doi.org/10.17605/OSF.IO/SNXQM.

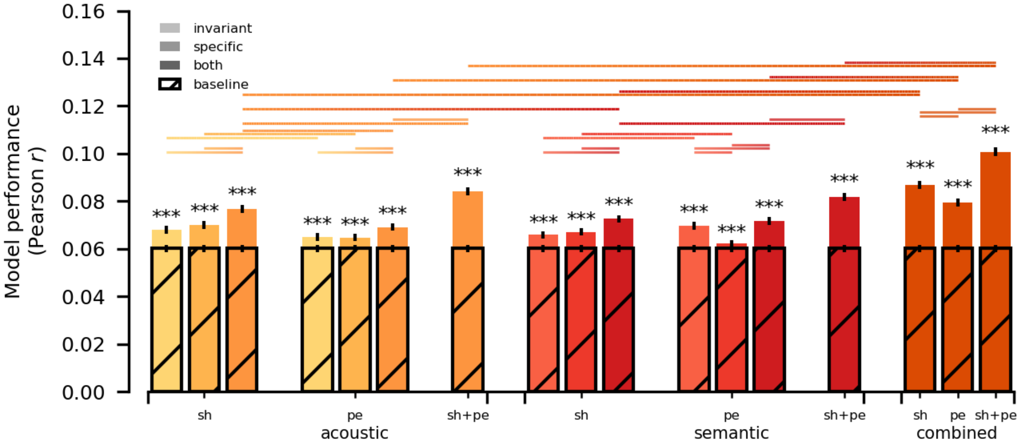


**Supplementary Fig. 7** Between-item RSA regression models using $k = 1$ predictions to validate results using a conventional approach and without spectrogram averaging (see Conventional representational similarity analysis). Results confirm the dominance of acoustic sharpening (sh) over acoustic prediction errors (pe), as well as the dominance of acoustic over semantic sharpening. Unlike the within-item cRSM regression, this approach also shows modest additive variance explained by both acoustic prediction errors and semantic sharpening and prediction errors, though variance uniquely attributable to either is comparatively smaller (see Supplementary Table 7). Therefore, this does not change our interpretation that acoustic sharpening is the dominant computation at the sensory level, but suggests that multiple computations may co-occur at the same level to varying degrees. Individual bars represent group means, with black lines indicating 95%-confidence intervals around the mean. ***, **, * $p \leq 1e^{-3}$, $p \leq 1e^{-2}$, and $p \leq 5e^{-2}$ for model comparisons against baseline, respectively. Big bars between models indicate $p \leq 5e^{-2}$. Data and code supporting these findings are available from https://doi.org/10.17605/OSF.IO/SNXQM.

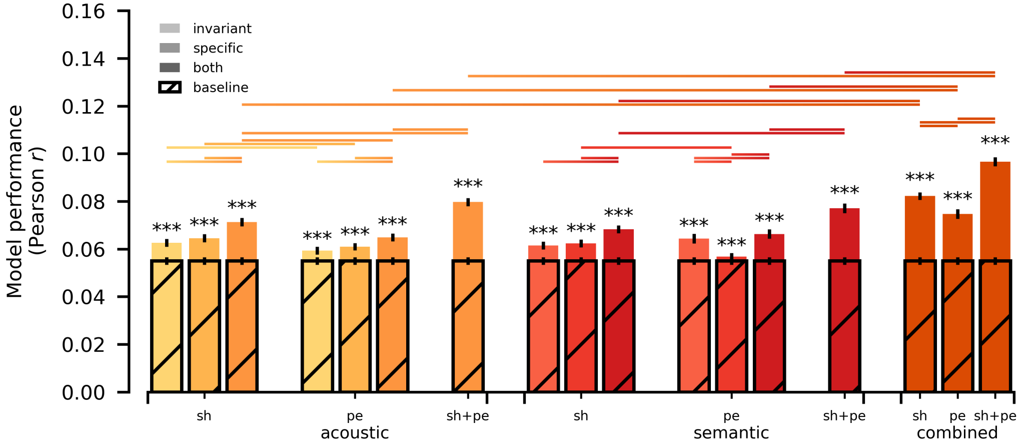


**Supplementary Fig. 8** Between-item RSA regression models, setting $k = 1$ to avoid averaging and restricting the temporal window of the analysis to the shortest possible prediction (0.0-0.48$s$) such that differences in length could not affect results (see Conventional representational similarity analysis). Results corroborate the pattern observed without temporal restrictions (see Supplementary Fig. 7), ruling out prediction length as a confounding factor (see Supplementary Table 8). Individual bars represent group means, with black lines indicating 95%-confidence intervals around the mean. ***, **, * indicate $p \leq 1e^{-3}$, $p \leq 1e^{-2}$, and $p \leq 5e^{-2}$, respectively. Big bars between groups indicate $p \leq 5e^{-2}$. Data and code supporting these findings are available from https://doi.org/10.17605/OSF.IO/SNXQM.

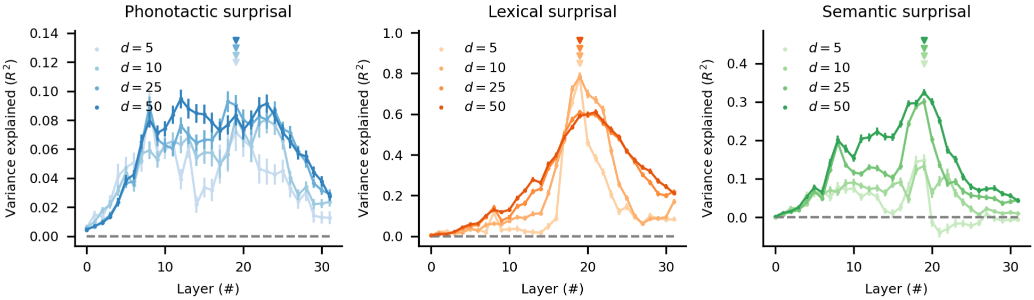


**Supplementary Fig. 9** To identify the layer within word2vec2.0 that best captured phonotactic, lexical and semantic surprisal, we performed back-to-back decoding[2] over an independent set of narrative stimuli (see Pretrained transformers as statistical surrogates). Here, we show the relative causal contribution of each feature across layers, for a number of PCA projections with dimensionality *d*. Small dots represent means, with 95%-confidence intervals around them. Downward-facing triangles represent the best layer overall for each *d*. Crucially, for all *d* tested, layer 19 (transformer layer 12) was identified as the most informative layer. This is in accordance with recent results finding that middle layers of transformers tend to reflect neural processing best[3]. Data and code supporting these findings are available from https://doi.org/10.17605/OSF.IO/SNXQM.

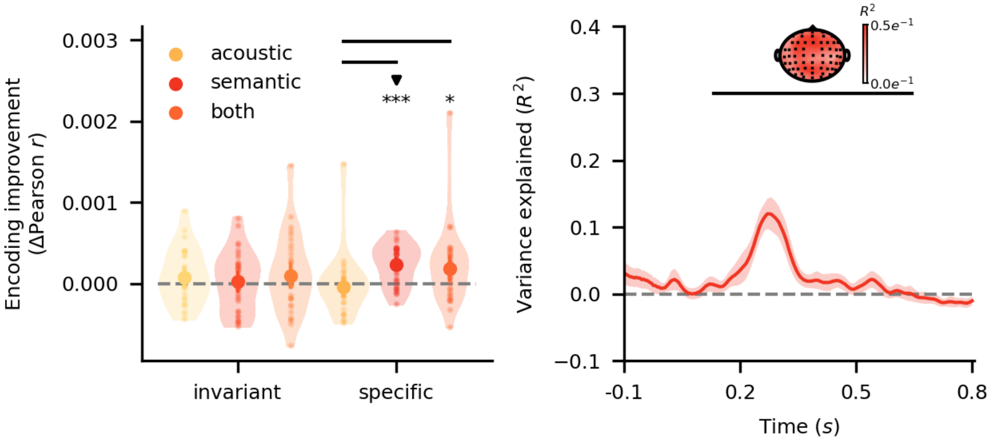


**Supplementary Fig. 10** To test the robustness of the effect of speaker-specific semantic surprisal, we refit the models using $d = 10$ features from transformer activations. Left: This revealed that, again, only speaker-specific semantic surprisal improved encoding performance—though here results varied more strongly, likely due to the now relatively high number of predictors within the model. Big dots represent group means, with 95%-confidence intervals around them. Small dots represent individual participants. ***, **, * indicate $p \leq 1e^{-3}$, $p \leq 1e^{-2}$, and $p \leq 5e^{-2}$, respectively. The downward-facing triangle marks the best model overall. Big bold lines indicate $p \leq 5e^{-2}$. Right: As before, speaker-specific semantic surprisal explained significant variance, with a cluster across a wide array of sensors around $110ms$-$400ms$. The inlaid topography shows variance explained at each sensor position within the cluster, with channels contributing to the cluster highlighted in black. Lines indicate means, with 95%-confidence intervals as shaded areas around them. Bold black lines indicate $p \leq 5e^{-2}$. Data and code supporting these findings are available from https://doi.org/10.17605/OSF.IO/SNXQM.

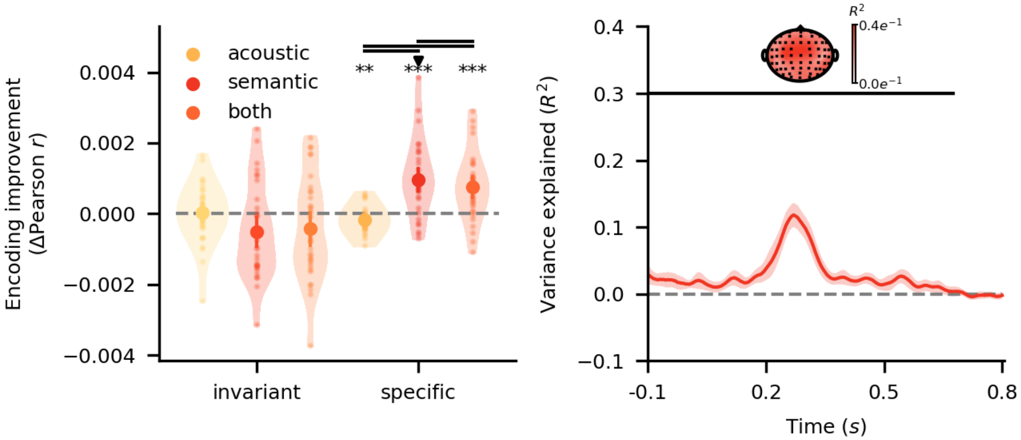


**Supplementary Fig. 11** To further probe the robustness of speaker-specific semantic surprisal, we refit the models using control predictors derived from the word that was more likely under the speaker prior. Consequently, these models were implicitly biassed against speaker-specific semantic surprisal, as this disambiguation of the morph meant some speaker-specific information was already encoded in the control models. Left: Again, we find that speaker-specific semantic surprisal was the only predictor that significantly improved model performance. Big dots represent group means, with 95%-confidence intervals around them. Small dots represent individual participants. ***, **, and * represent $p \leq 1e^{-3}$, $p \leq 1e^{-2}$, and $p \leq 5e^{-2}$, respectively. The downward-facing triangle marks the best model overall. Big black lines indicate $p \leq 5e^{-2}$. Right: Again, we find that speaker-specific semantic surprisal explains significant variance, with a cluster across all sensors between $-100ms$-$410ms$. The inlaid topography shows variance explained at each sensor position, with sensors contributing to the cluster marked in black. Lines indicate means, with 95%-confidence intervals around them. Big bold lines indicate $p \leq 5e^{-2}$. Data and code supporting these findings are available from https://doi.org/10.17605/OSF.IO/SNXQM.

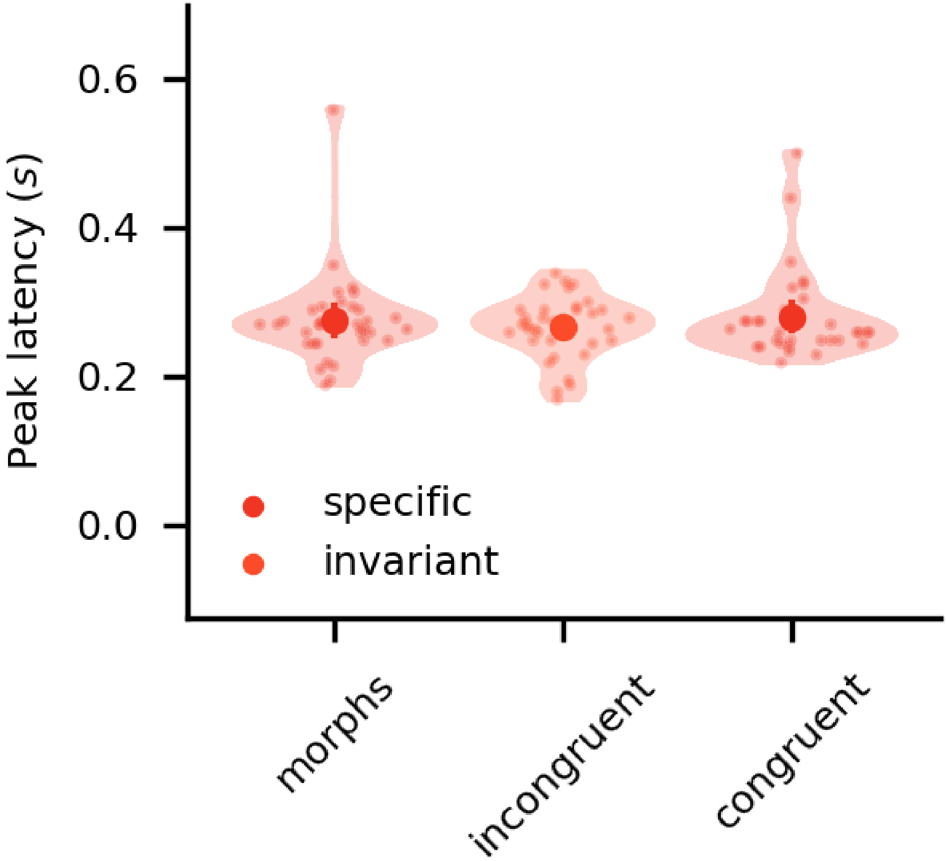


**Supplementary Fig. 12** Peak latencies of semantic surprisal effects across morph, congruent and incongruent trials, demonstrating that there was no significant difference between peak latencies. Note that these represent raw latency estimates over the temporal extent of the cluster. No jackknife procedure was applied. Big dots indicate group means, with 95%-confidence intervals around them. Small dots represent individual participants. Data and code supporting these findings are available from https://doi.org/10.17605/OSF.IO/SNXQM.

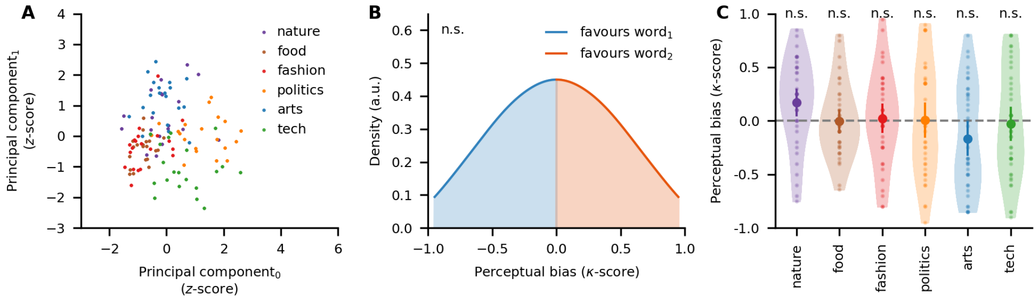


**Supplementary Fig. 13** Overview of stimulus materials. **A** Visualisation of the semantic space spanned by the words used in this experiment, as projected into two dimensions using PCA. Dots represent individual words, coloured by the semantic context they were most likely under. **B** Gaussian approximation of the distribution of perceptual bias of morphs across the experiment, showing that stimuli exhibit no systematic bias. **C** Perceptual bias of morphs for individual contexts. Big dots indicate context means, with 95%-confidence intervals around them. Small dots represent individual morphs. Again, no systematic biases exist. Data and code supporting these findings are available from https://doi.org/10.17605/OSF.IO/SNXQM.

#### 2 Supplementary Tables

| Coefficient | Estimate | Std. Error | $z$-value | $p$-value |
|---|---|---|---|---|
| (Intercept) | 0.27 | 0.06 | 4.42 | 9.824392e-06 |
| $t$ | -0.04 | 0.03 | -1.47 | 1.413237e-01 |
| $\kappa$ | -0.62 | 0.11 | -5.81 | 6.261726e-09 |
| fit | 1.64 | 0.12 | 13.11 | 2.712021e-39 |
| $t \times \kappa$ | 0.39 | 0.06 | 6.09 | 1.135522e-09 |
| $t\times$fit | 0.43 | 0.03 | 12.77 | 2.405336e-37 |

**Supplementary Table 1.** Results from generalised linear model over responses from online experiment.

| Coefficient | Estimate | Std. Error | $z$-value | $p$-value |
|---|---|---|---|---|
| (Intercept) | 0.13 | 0.05 | 2.84 | 4.546977e-03 |
| $t$ | -0.26 | 0.03 | -8.00 | 1.197850e-15 |
| $\kappa$ | -0.14 | 0.09 | -1.60 | 1.102186e-01 |
| fit | 1.35 | 0.06 | 20.99 | 8.269810e-98 |
| $t \times \kappa$ | 0.42 | 0.06 | 6.62 | 3.606962e-11 |
| $t\times$ fit | 0.38 | 0.03 | 11.07 | 1.811167e-28 |

**Supplementary Table 2.** Results from generalised linear model over responses from EEG experiment.

| Frequency band | M | Std. Dev. | df | $t$-value | $p$-value |
|---|---|---|---|---|---|
| 0 | 0.06 | 0.03 | 34 | 10.56 | 3.151979e-11 |
| 1 | 0.06 | 0.03 | 34 | 11.03 | 1.352901e-11 |
| 2 | 0.06 | 0.04 | 34 | 9.30 | 5.005196e-10 |
| 3 | 0.06 | 0.04 | 34 | 9.45 | 3.847424e-10 |
| 4 | 0.06 | 0.04 | 34 | 10.13 | 7.584404e-11 |
| 5 | 0.07 | 0.03 | 34 | 12.00 | 1.983854e-12 |
| 6 | 0.08 | 0.03 | 34 | 13.22 | 1.423830e-13 |
| 7 | 0.08 | 0.04 | 34 | 12.86 | 2.992608e-13 |
| 8 | 0.09 | 0.04 | 34 | 14.33 | 1.534314e-14 |
| 9 | 0.09 | 0.04 | 34 | 13.60 | 6.844391e-14 |
| 10 | 0.08 | 0.04 | 34 | 11.84 | 2.616005e-12 |
| 11 | 0.07 | 0.04 | 34 | 10.90 | 1.581847e-11 |
| 12 | 0.06 | 0.03 | 34 | 10.69 | 2.490932e-11 |
| 13 | 0.06 | 0.03 | 34 | 11.21 | 9.356256e-12 |
| 14 | 0.07 | 0.03 | 34 | 11.35 | 7.484068e-12 |
| 15 | 0.07 | 0.04 | 34 | 11.34 | 7.129978e-12 |
| 16 | 0.06 | 0.03 | 34 | 11.00 | 1.357066e-11 |
| 17 | 0.07 | 0.03 | 34 | 11.36 | 7.745446e-12 |
| 18 | 0.06 | 0.03 | 34 | 11.99 | 1.924524e-12 |
| 19 | 0.05 | 0.03 | 34 | 10.25 | 6.210367e-11 |
| 20 | 0.03 | 0.02 | 34 | 7.44 | 6.303351e-08 |
| 21 | 0.03 | 0.03 | 34 | 5.87 | 2.529152e-06 |
| 22 | 0.03 | 0.03 | 34 | 6.01 | 2.514777e-06 |
| 23 | 0.03 | 0.03 | 34 | 5.74 | 1.852334e-06 |
| 24 | 0.03 | 0.03 | 34 | 6.87 | 2.638222e-07 |
| 25 | 0.04 | 0.02 | 34 | 8.32 | 6.177527e-09 |
| 26 | 0.06 | 0.02 | 34 | 13.36 | 1.084790e-13 |
| 27 | 0.10 | 0.04 | 34 | 15.24 | 2.594835e-15 |

**Supplementary Table 3.** Results from stimulus reconstructions for each frequency band.

| contrast | M | Std. Dev | df | $t$-value | $p$-value |
|---|---|---|---|---|---|
| 3-1 | 0.092 | 0.011 | 34 | 49.284 | 2.954380e-32 |
| 5-3 | 0.034 | 0.007 | 34 | 27.689 | 5.264768e-24 |
| 7-5 | 0.014 | 0.003 | 34 | 24.428 | 2.674880e-22 |
| 9-7 | 0.008 | 0.004 | 34 | 13.033 | 5.308751e-14 |
| 11-9 | 0.004 | 0.003 | 34 | 9.122 | 5.810556e-10 |
| 13-11 | 0.004 | 0.003 | 34 | 8.287 | 4.540814e-09 |
| 15-13 | 0.003 | 0.002 | 34 | 7.560 | 2.655277e-08 |
| 17-15 | 0.001 | 0.002 | 34 | 3.150 | 3.398163e-03 |
| 19-17 | 0.003 | 0.002 | 34 | 6.885 | 1.250519e-07 |

**Supplementary Table 4.** Results from consecutive contrasts of top-$k$ within-item cRSM regression models.

| contrast | M | Std. Dev | df | $t$-value | $p$-value |
|---|---|---|---|---|---|
| ac.inv.-baseline | 0.0032 | 0.0014 | 34 | 13.1557 | 1.083377e-13 |
| ac.spc.-baseline | 0.0066 | 0.0013 | 34 | 29.3987 | 2.415507e-24 |
| ac.spc.-ac.inv. | 0.0034 | 0.0014 | 34 | 14.1754 | 1.331002e-14 |
| ac.bth.-baseline | 0.0107 | 0.0017 | 34 | 36.9431 | 1.487629e-27 |
| ac.bth.-ac.inv. | 0.0075 | 0.0015 | 34 | 29.7752 | 1.714360e-24 |
| ac.bth.-ac.spc. | 0.0041 | 0.0011 | 34 | 20.7433 | 1.481206e-19 |
| sem.inv.-baseline | -0.0001 | 0.0004 | 34 | -0.8850 | 1.000000e+00 |
| sem.spc.-baseline | -0.0004 | 0.0002 | 34 | -8.3291 | 1.312180e-08 |
| sem.spc.-sem.inv. | -0.0003 | 0.0004 | 34 | -4.4158 | 8.721333e-04 |
| sem.bth.-baseline | -0.0005 | 0.0004 | 34 | -6.8839 | 7.533303e-07 |
| sem.bth.-sem.inv. | -0.0005 | 0.0001 | 34 | -22.5252 | 1.133089e-20 |
| sem.bth.-sem.spc. | -0.0002 | 0.0004 | 34 | -2.5786 | 1.153901e-01 |
| sem.inv.-ac.inv. | -0.0031 | 0.0032 | 34 | -5.6651 | 2.346169e-05 |
| sem.spc.-ac.spc. | -0.0068 | 0.0026 | 34 | -15.5283 | 9.511598e-16 |
| sem.bth.-ac.bth. | -0.0111 | 0.0028 | 34 | -23.0672 | 5.779476e-21 |

**Supplementary Table 5.** Results from contrasts in within-item cRSM regression models using $k = 19$.

| contrast | M | Std. Dev | df | $t$-value | $p$-value |
|---|---|---|---|---|---|
| ac.inv.: true-foil | 0.002073 | 0.007127 | 34 | 1.696327 | 3.958520e-01 |
| ac.spc: true-foil | 0.007507 | 0.006593 | 34 | 6.638746 | 7.745975e-07 |
| ac.both: true-foil | 0.007945 | 0.007985 | 34 | 5.801883 | 7.787670e-06 |
| sem.inv.: true-foil | -0.000825 | 0.006090 | 34 | -0.790213 | 1.000000e+00 |
| sem.spc: true-foil | -0.000710 | 0.006138 | 34 | -0.674545 | 5.045258e-01 |
| sem.both: true-foil | -0.000769 | 0.006059 | 34 | -0.739759 | 9.290471e-01 |

**Supplementary Table 6.** Results from within-item cRSM regression models, comparing foil priors to real participant-specific priors (see Validating top-$k$ acoustic predictions).

| contrast | M | Std. Dev | df | $t$-value | $p$-value |
|---|---|---|---|---|---|
| baseline-acc.inv. (sh) | -0.008 | 0.004 | 34 | -12.56 | 6.8e-13 |
| baseline-acc.spc. (sh) | -0.010 | 0.005 | 34 | -12.14 | 1.6e-12 |
| baseline-acc.bth. (sh) | -0.017 | 0.006 | 34 | -17.09 | 1.2e-16 |
| baseline-acc.inv. (pe) | -0.005 | 0.002 | 34 | -12.85 | 3.9e-13 |
| baseline-acc.spc. (pe) | -0.004 | 0.002 | 34 | -14.34 | 2.0e-14 |
| baseline-acc.bth. (pe) | -0.009 | 0.003 | 34 | -20.08 | 8.6e-19 |
| baseline-acc.bth. (sh+pe) | -0.024 | 0.006 | 34 | -24.36 | 2.1e-21 |
| baseline-sem.inv. (sh) | -0.006 | 0.003 | 34 | -13.01 | 2.9e-13 |
| baseline-sem.spc. (sh) | -0.007 | 0.005 | 34 | -8.10 | 3.0e-08 |
| baseline-sem.bth. (sh) | -0.012 | 0.005 | 34 | -13.95 | 4.2e-14 |
| baseline-sem.inv. (pe) | -0.010 | 0.005 | 34 | -12.24 | 1.3e-12 |
| baseline-sem.spc. (pe) | -0.002 | 0.002 | 34 | -5.63 | 2.6e-05 |
| baseline-sem.bth. (pe) | -0.012 | 0.005 | 34 | -12.86 | 3.9e-13 |
| baseline-sem.bth. (sh+pe) | -0.022 | 0.007 | 34 | -17.05 | 1.3e-16 |
| baseline-acc.sem. (sh) | -0.027 | 0.007 | 34 | -23.14 | 1.1e-20 |
| baseline-acc.sem. (pe) | -0.019 | 0.005 | 34 | -21.90 | 5.8e-20 |
| baseline-acc.sem. (sh+pe) | -0.041 | 0.008 | 34 | -29.33 | 5.1e-24 |
| acc.inv. (sh)-acc.spc. (sh) | -0.002 | 0.006 | 34 | -2.08 | 2.2e-01 |
| acc.inv. (sh)-acc.bth. (sh) | -0.009 | 0.004 | 34 | -11.49 | 6.2e-12 |
| acc.spc. (sh)-acc.bth. (sh) | -0.007 | 0.003 | 34 | -12.04 | 1.9e-12 |
| acc.inv. (pe)-acc.spc. (pe) | 0.000 | 0.003 | 34 | 0.55 | 5.9e-01 |
| acc.inv. (pe)-acc.bth. (pe) | -0.004 | 0.002 | 34 | -13.64 | 7.9e-14 |
| acc.spc. (pe)-acc.bth. (pe) | -0.004 | 0.002 | 34 | -11.90 | 2.5e-12 |
| acc.inv. (sh)-acc.inv. (pe) | 0.003 | 0.005 | 34 | 3.71 | 5.9e-03 |
| acc.spc. (sh)-acc.spc. (pe) | 0.005 | 0.004 | 34 | 7.04 | 5.6e-07 |
| acc.bth. (sh)-acc.bth. (pe) | 0.008 | 0.006 | 34 | 7.46 | 1.8e-07 |
| acc.bth. (sh)-acc.bth. (sh+pe) | -0.007 | 0.002 | 34 | -20.06 | 8.7e-19 |
| acc.bth. (pe)-acc.bth. (sh+pe) | -0.015 | 0.005 | 34 | -16.47 | 3.5e-16 |
| sem.inv. (sh)-sem.spc. (sh) | -0.001 | 0.005 | 34 | -1.43 | 4.9e-01 |
| sem.inv. (sh)-sem.bth. (sh) | -0.007 | 0.004 | 34 | -8.81 | 4.6e-09 |
| sem.spc. (sh)-sem.bth. (sh) | -0.005 | 0.003 | 34 | -10.93 | 2.2e-11 |
| sem.inv. (pe)-sem.spc. (pe) | 0.008 | 0.004 | 34 | 10.71 | 3.5e-11 |
| sem.inv. (pe)-sem.bth. (pe) | -0.002 | 0.002 | 34 | -6.97 | 6.3e-07 |
| sem.spc. (pe)-sem.bth. (pe) | -0.010 | 0.005 | 34 | -11.39 | 7.6e-12 |
| sem.inv. (sh)-sem.inv. (pe) | -0.004 | 0.005 | 34 | -4.73 | 3.4e-04 |
| sem.spc. (sh)-sem.spc. (pe) | 0.005 | 0.005 | 34 | 5.99 | 9.6e-06 |
| sem.bth. (sh)-sem.bth. (pe) | 0.001 | 0.006 | 34 | 0.85 | 8.1e-01 |
| sem.bth. (sh)-sem.bth. (sh+pe) | -0.009 | 0.004 | 34 | -12.10 | 1.7e-12 |
| sem.bth. (pe)-sem.bth. (sh+pe) | -0.010 | 0.004 | 34 | -14.14 | 3.0e-14 |
| acc.bth. (sh)-sem.bth. (sh) | 0.004 | 0.008 | 34 | 3.23 | 1.9e-02 |
| acc.bth. (pe)-sem.bth. (pe) | -0.003 | 0.006 | 34 | -2.62 | 7.7e-02 |

| contrast | M | Std. Dev | df | $t$-value | $p$-value |
|---|---|---|---|---|---|
| acc.bth. (sh+pe)-sem.bth. (sh+pe) | 0.002 | 0.009 | 34 | 1.45 | 6.3e-01 |
| acc.bth. (sh)-acc.sem. (sh) | -0.010 | 0.004 | 34 | -13.37 | 1.4e-13 |
| sem.bth. (sh)-acc.sem. (sh) | -0.014 | 0.005 | 34 | -16.29 | 4.8e-16 |
| acc.bth. (pe)-acc.sem. (pe) | -0.010 | 0.005 | 34 | -12.70 | 5.1e-13 |
| sem.bth. (pe)-acc.sem. (pe) | -0.008 | 0.002 | 34 | -18.21 | 1.7e-17 |
| acc.sem. (sh)-acc.sem. (pe) | 0.007 | 0.006 | 34 | 6.75 | 1.1e-06 |
| acc.sem. (sh)-acc.sem. (sh+pe) | -0.014 | 0.004 | 34 | -21.38 | 1.2e-19 |
| acc.sem. (pe)-acc.sem. (sh+pe) | -0.021 | 0.005 | 34 | -23.88 | 3.9e-21 |
| acc.bth. (sh+pe)-acc.sem. (sh+pe) | -0.017 | 0.006 | 34 | -15.63 | 1.6e-15 |
| sem.bth. (sh+pe)-acc.sem. (sh+pe) | -0.019 | 0.005 | 34 | -23.07 | 1.1e-20 |

**Supplementary Table 7.** Results from between-item RSA regression models using $k = 1$ (see Conventional representational similarity analysis).

| contrast | M | Std. Dev | df | $t$-value | $p$-value |
|---|---|---|---|---|---|
| baseline-acc.inv. (sh) | -0.008 | 0.004 | 34 | -11.22 | 1.7e-11 |
| baseline-acc.spc. (sh) | -0.010 | 0.004 | 34 | -13.44 | 1.3e-13 |
| baseline-acc.bth. (sh) | -0.016 | 0.005 | 34 | -20.10 | 8.8e-19 |
| baseline-acc.inv. (pe) | -0.004 | 0.002 | 34 | -10.74 | 5.3e-11 |
| baseline-acc.spc. (pe) | -0.006 | 0.004 | 34 | -9.82 | 4.9e-10 |
| baseline-acc.bth. (pe) | -0.010 | 0.004 | 34 | -16.20 | 5.9e-16 |
| baseline-acc.bth. (sh+pe) | -0.025 | 0.005 | 34 | -26.22 | 1.9e-22 |
| baseline-sem.inv. (sh) | -0.006 | 0.003 | 34 | -11.09 | 2.3e-11 |
| baseline-sem.spc. (sh) | -0.007 | 0.006 | 34 | -7.60 | 1.3e-07 |
| baseline-sem.bth. (sh) | -0.013 | 0.006 | 34 | -13.60 | 9.6e-14 |
| baseline-sem.inv. (pe) | -0.009 | 0.006 | 34 | -8.41 | 1.5e-08 |
| baseline-sem.spc. (pe) | -0.002 | 0.002 | 34 | -4.50 | 6.9e-04 |
| baseline-sem.bth. (pe) | -0.011 | 0.007 | 34 | -9.31 | 1.7e-09 |
| baseline-sem.bth. (sh+pe) | -0.022 | 0.008 | 34 | -15.29 | 3.3e-15 |
| baseline-acc.sem. (sh) | -0.027 | 0.005 | 34 | -29.14 | 6.2e-24 |
| baseline-acc.sem. (pe) | -0.020 | 0.007 | 34 | -16.54 | 3.3e-16 |
| baseline-acc.sem. (sh+pe) | -0.042 | 0.007 | 34 | -32.46 | 1.8e-25 |
| acc.inv. (sh)-acc.spc. (sh) | -0.002 | 0.006 | 34 | -1.77 | 4.2e-01 |
| acc.inv. (sh)-acc.bth. (sh) | -0.009 | 0.004 | 34 | -12.61 | 7.7e-13 |
| acc.spc. (sh)-acc.bth. (sh) | -0.007 | 0.004 | 34 | -10.55 | 8.1e-11 |
| acc.inv. (pe)-acc.spc. (pe) | -0.002 | 0.005 | 34 | -2.00 | 3.2e-01 |
| acc.inv. (pe)-acc.bth. (pe) | -0.006 | 0.003 | 34 | -9.33 | 1.7e-09 |
| acc.spc. (pe)-acc.bth. (pe) | -0.004 | 0.002 | 34 | -10.15 | 2.1e-10 |
| acc.inv. (sh)-acc.inv. (pe) | 0.003 | 0.004 | 34 | 4.64 | 5.5e-04 |
| acc.spc. (sh)-acc.spc. (pe) | 0.004 | 0.005 | 34 | 4.59 | 5.9e-04 |
| acc.bth. (sh)-acc.bth. (pe) | 0.006 | 0.005 | 34 | 7.47 | 1.7e-07 |
| acc.bth. (sh)-acc.bth. (sh+pe) | -0.008 | 0.003 | 34 | -16.65 | 2.8e-16 |
| acc.bth. (pe)-acc.bth. (sh+pe) | -0.015 | 0.004 | 34 | -20.00 | 1.0e-18 |
| sem.inv. (sh)-sem.spc. (sh) | -0.001 | 0.006 | 34 | -0.88 | 3.9e-01 |
| sem.inv. (sh)-sem.bth. (sh) | -0.007 | 0.005 | 34 | -8.74 | 6.5e-09 |
| sem.spc. (sh)-sem.bth. (sh) | -0.006 | 0.004 | 34 | -8.99 | 3.4e-09 |
| sem.inv. (pe)-sem.spc. (pe) | 0.008 | 0.006 | 34 | 7.62 | 1.3e-07 |
| sem.inv. (pe)-sem.bth. (pe) | -0.002 | 0.002 | 34 | -6.28 | 5.2e-06 |
| sem.spc. (pe)-sem.bth. (pe) | -0.010 | 0.007 | 34 | -8.43 | 1.5e-08 |
| sem.inv. (sh)-sem.inv. (pe) | -0.003 | 0.007 | 34 | -2.55 | 1.2e-01 |
| sem.spc. (sh)-sem.spc. (pe) | 0.006 | 0.006 | 34 | 5.37 | 6.8e-05 |
| sem.bth. (sh)-sem.bth. (pe) | 0.002 | 0.008 | 34 | 1.48 | 5.9e-01 |
| sem.bth. (sh)-sem.bth. (sh+pe) | -0.009 | 0.006 | 34 | -9.20 | 2.2e-09 |
| sem.bth. (pe)-sem.bth. (sh+pe) | -0.011 | 0.005 | 34 | -12.52 | 9.1e-13 |

| contrast | M | Std. Dev | df | $t$-value | $p$-value |
|---|---|---|---|---|---|
| acc.bth. (sh)-sem.bth. (sh) | 0.003 | 0.008 | 34 | 2.08 | 3.2e-01 |
| acc.bth. (pe)-sem.bth. (pe) | -0.001 | 0.008 | 34 | -1.03 | 6.2e-01 |
| acc.bth. (sh+pe)-sem.bth. (sh+pe) | 0.003 | 0.011 | 34 | 1.40 | 5.1e-01 |
| acc.bth. (sh)-acc.sem. (sh) | -0.011 | 0.005 | 34 | -12.39 | 1.2e-12 |
| sem.bth. (sh)-acc.sem. (sh) | -0.014 | 0.005 | 34 | -17.50 | 6.3e-17 |
| acc.bth. (pe)-acc.sem. (pe) | -0.010 | 0.006 | 34 | -9.06 | 3.0e-09 |
| sem.bth. (pe)-acc.sem. (pe) | -0.008 | 0.003 | 34 | -15.04 | 5.3e-15 |
| acc.sem. (sh)-acc.sem. (pe) | 0.008 | 0.008 | 34 | 5.59 | 3.8e-05 |
| acc.sem. (sh)-acc.sem. (sh+pe) | -0.014 | 0.005 | 34 | -16.43 | 4.0e-16 |
| acc.sem. (pe)-acc.sem. (sh+pe) | -0.022 | 0.005 | 34 | -26.12 | 2.1e-22 |
| acc.bth. (sh+pe)-acc.sem. (sh+pe) | -0.017 | 0.007 | 34 | -13.38 | 1.5e-13 |
| sem.bth. (sh+pe)-acc.sem. (sh+pe) | -0.020 | 0.005 | 34 | -22.69 | 1.9e-20 |

**Supplementary Table 8.** Results from between-item RSA regression models using $k = 1$ and restricting the temporal window to 0.0-0.48$s$ to rule out prediction length as a confounding factor (see Conventional representational similarity analysis).

| contrast | M | Std. Dev. | df | $t$-value | $p$-value |
|---|---|---|---|---|---|
| inv. ac.-baseline | 0.000076 | 0.000316 | 34 | 1.399752 | 1.000000 |
| inv. sem.-baseline | 0.000023 | 0.000350 | 34 | 0.376861 | 0.708619 |
| inv. sem.-ac. | -0.000053 | 0.000508 | 34 | -0.611232 | 1.000000 |
| inv. bth.-baseline | 0.000099 | 0.000438 | 34 | 1.311878 | 0.991752 |
| inv. bth.-ac. | 0.000023 | 0.000333 | 34 | 0.397016 | 1.000000 |
| inv. bth.-sem. | 0.000076 | 0.000352 | 34 | 1.258929 | 0.866525 |
| spc. ac.-baseline | -0.000040 | 0.000329 | 34 | -0.707456 | 0.968209 |
| spc. sem.-baseline | 0.000236 | 0.000230 | 34 | 5.982584 | 0.000005 |
| spc. sem.-ac. | 0.000276 | 0.000428 | 34 | 3.754684 | 0.002601 |
| spc. bth.-baseline | 0.000188 | 0.000431 | 34 | 2.545933 | 0.046801 |
| spc. bth.-ac. | 0.000228 | 0.000235 | 34 | 5.660051 | 0.000012 |
| spc. bth.-sem. | -0.000048 | 0.000401 | 34 | -0.696623 | 0.490774 |

**Supplementary Table 9.** Results from contrasts in single-trial encoding models using pretrained transformers with $d = 10$.

| contrast | M | Std. Dev. | df | $t$-value | $p$-value |
|---|---|---|---|---|---|
| inv. ac.-baseline | 0.000026 | 0.000799 | 34 | 0.191701 | 0.849117 |
| inv. sem.-baseline | -0.000517 | 0.001237 | 34 | -2.439485 | 0.080313 |
| inv. sem.-ac. | -0.000544 | 0.001274 | 34 | -2.488849 | 0.107237 |
| inv. bth.-baseline | -0.000431 | 0.001368 | 34 | -1.838238 | 0.224332 |
| inv. bth.-ac. | -0.000458 | 0.001081 | 34 | -2.469447 | 0.093559 |
| inv. bth.-sem. | 0.000086 | 0.000711 | 34 | 0.705434 | 0.970692 |
| spc. ac.-baseline | -0.000175 | 0.000327 | 34 | -3.116407 | 0.003709 |
| spc. sem.-baseline | 0.000961 | 0.000985 | 34 | 5.687480 | 0.000009 |
| spc. sem.-ac. | 0.001136 | 0.001106 | 34 | 5.989475 | 0.000005 |
| spc. bth.-baseline | 0.000760 | 0.000896 | 34 | 4.947531 | 0.000060 |
| spc. bth.-ac. | 0.000935 | 0.000943 | 34 | 5.778516 | 0.000008 |
| spc. bth.-sem. | -0.000201 | 0.000314 | 34 | -3.725773 | 0.001410 |

**Supplementary Table 10.** Results from contrasts in single-trial encoding models using control predictors derived from target words.

| contrast | M | Std. Dev. | df | *t*-value | *p*-value |
|---|---|---|---|---|---|
| inc. inv. ac.-baseline | -0.000315 | 0.001721 | 34 | -1.065444 | 5.883657e-01 |
| inc. inv. sem.-baseline | 0.001820 | 0.003506 | 34 | 3.026296 | 1.877703e-02 |
| inc. inv. sem.-ac. | 0.002134 | 0.003596 | 34 | 3.460802 | 8.825596e-03 |
| inc. inv. bth.-baseline | 0.001554 | 0.003933 | 34 | 2.304360 | 8.231825e-02 |
| inc. inv. bth.-ac. | 0.001869 | 0.003298 | 34 | 3.304056 | 1.125702e-02 |
| inc. inv. bth.-sem. | -0.000265 | 0.001571 | 34 | -0.984436 | 3.318550e-01 |
| inc. spc. ac.-baseline | -0.001189 | 0.000724 | 34 | -9.580910 | 2.070851e-10 |
| inc. spc. sem.-baseline | -0.001074 | 0.002560 | 34 | -2.446197 | 5.929342e-02 |
| inc. spc. sem.-ac. | 0.000116 | 0.002711 | 34 | 0.248509 | 8.052359e-01 |
| inc. spc. bth.-baseline | -0.002037 | 0.002557 | 34 | -4.646061 | 1.967241e-04 |
| inc. spc. bth.-ac. | -0.000848 | 0.002522 | 34 | -1.959992 | 1.164685e-01 |
| inc. spc. bth.-sem. | -0.000963 | 0.000794 | 34 | -7.071817 | 1.812218e-07 |
| con. inv. ac.-baseline | -0.000387 | 0.001380 | 34 | -1.634498 | 2.227622e-01 |
| con. inv. sem.-baseline | 0.001020 | 0.003020 | 34 | 1.969295 | 2.284476e-01 |
| con. inv. sem.-ac. | 0.001407 | 0.003301 | 34 | 2.484502 | 9.028907e-02 |
| con. inv. bth.-baseline | 0.000909 | 0.003185 | 34 | 1.663600 | 3.161534e-01 |
| con. inv. bth.-ac. | 0.001295 | 0.002896 | 34 | 2.608023 | 8.059220e-02 |
| con. inv. bth.-sem. | -0.000111 | 0.001388 | 34 | -0.467179 | 6.433524e-01 |
| con. spc. ac.-baseline | -0.000813 | 0.000919 | 34 | -5.157861 | 3.215765e-05 |
| con. spc. sem.-baseline | 0.002554 | 0.001775 | 34 | 8.392109 | 3.390330e-09 |
| con. spc. sem.-ac. | 0.003368 | 0.001905 | 34 | 10.309204 | 3.190655e-11 |
| con. spc. bth.-baseline | 0.001811 | 0.002108 | 34 | 5.007559 | 3.358951e-05 |
| con. spc. bth.-ac. | 0.002624 | 0.001774 | 34 | 8.622222 | 2.248759e-09 |
| con. spc. bth.-sem. | -0.000744 | 0.000999 | 34 | -4.342367 | 1.201464e-04 |

**Supplementary Table 11.** Results from contrasts in single-trial encoding models in the follow-up task.

| pair | contexts |
|---|---|
| Karte-Kette | tech-fashion |
| Anzüge-einzige | fashion-politics |
| Justiz-Notiz | politics-arts |
| Benutzer-Beschützer | tech-arts |
| Ballade-Panade | arts-food |
| Echtzeit-Bescheid | tech-politics |
| kürzen-würzen | fashion-food |
| Fährte-Konzerte | nature-arts |
| autokratisch-automatisch | politics-tech |
| Boot-Bit | nature-tech |
| Verwaltung-Vergeltung | politics-arts |
| kaputte-Kapuze | tech-fashion |
| Batterie-Fantasie | tech-arts |
| Brei-Schrei | food-arts |
| Agentur-Tastatur | politics-tech |
| segeln-Regeln | nature-politics |
| Olive-Motive | food-arts |
| Hitze-Witze | food-arts |
| Cloud-Braut | tech-fashion |
| Nacht-Recht | arts-politics |
| dörren-dürren | food-fashion |
| Brüste-Küste | fashion-nature |
| Regime-intim | politics-arts |
| Berg-Wert | nature-politics |
| Bühnen-grünen | arts-politics |
| schwimmen-Stimmen | nature-politics |
| Laser-Blazer | tech-fashion |
| schuften-duften | nature-food |
| Nadeln-radeln | fashion-nature |
| Wellen-Zellen | nature-tech |
| Platine-Gardine | tech-fashion |
| Tee-See | food-nature |
| betrogen-Beethoven | politics-arts |
| Münze-Künste | politics-arts |
| satt-Watt | food-tech |
| Professor-Prozessor | politics-tech |
| Stiefel-Staffel | fashion-arts |
| braune-Pflaume | fashion-food |
| Kopf-Topf | fashion-food |
| erkunden-erfunden | nature-arts |
| Zutaten-Flughafen | food-nature |

| pair | contexts |
|---|---|
| Lichter-Dichter | nature-arts |
| lecker-Hacker | food-tech |
| Daten-waten | tech-nature |
| Schal-Saal | fashion-arts |
| Feld-Geld | nature-politics |
| Schaum-Saum | food-fashion |
| Treibsand-Breitband | nature-tech |
| Meinungen-Leitungen | politics-tech |
| Mais-weiß | food-fashion |
| Kragen-fragen | fashion-politics |
| Hemd-Held | fashion-arts |
| suchen-Kuchen | nature-food |
| Spitzel-Pixel | politics-tech |
| Kopftuch-Kochbuch | fashion-food |
| Kayak-Cognac | nature-food |
| Wälder-Wähler | nature-politics |
| Volt-Wild | tech-nature |
| Schurke-Gurke | arts-food |
| Schmuck-Schluck | fashion-food |

**Supplementary Table 12.** All stimulus pairs along with their corresponding context pairs used in the present study.

### 3 Full list of legends

**Supplementary Fig. 1** Summary of key behavioural results from the online experiment one (top row) and EEG experiment two (bottom row). In both experiments, participants initially relied on remaining acoustic properties (i.e., morph bias to one of the words within each word pair), but decreased this reliance over time (left). Participants increasingly relied on the probability of the word given the speaker instead (right). Lines represent means, with shaded areas representing 95%-confidence intervals. For details, see Supplementary Table 1-2. Data and code supporting these findings are available from https://doi.org/10.17605/OSF.IO/SNXQM.

**Supplementary Fig. 2** Accuracy of stimulus reconstruction models[1] in all 28 individual frequency bands. Bold dots indicate means, with 95%-confidence intervals around them. Small dots represent individual participants. Inlaid topographies show the decoded pattern for this frequency band. ***, **, * represent $p \leq 1e^{-3}$, $p \leq 1e^{-2}$ and $p \leq 5e^{-2}$, respectively. For details, see Supplementary Table 3. Data and code supporting these findings are available from https://doi.org/10.17605/OSF.IO/SNXQM.

**Supplementary Fig. 3** We systematically refit models with increasing $k$ to test whether the brain makes a specific number of predictions $k$. Big dots represent group means, with 95%-confidence intervals around them. Small dots indicate individual participants. Big black bars on top indicate $p \leq 5e^{-2}$ for any contrast. For more details, see Supplementary Table 4. Data and code supporting these findings are available from https://doi.org/10.17605/OSF.IO/SNXQM.

**Supplementary Fig. 4** Similarity encoding models using $k = 19$ predictions, because increasing the number of predictions $k$ yielded significant improvements in performance. Left: Results show that both speaker-invariant and speaker-specific acoustic predictions improve model performance, with the best model incorporating both at once. Critically, purely semantic predictions failed to improve performance. This is in line with our results based on $k = 5$, reported in Fig. 2. Big dots represent group means, with 95%-confidence intervals around them. Small dots represent individual participants. ***, **, * indicate $p \leq 1e^{-3}$, $p \leq 1e^{-2}$, and $p \leq 5e^{-2}$, respectively. The downward-facing triangle marks the best model overall. Big black bars between groups indicate $p \leq 5e^{-2}$. Right: Coefficients for speaker-specific acoustic predictions showed, again, that there was significant sharpening of neural representations, with a cluster between $0ms$-$1000ms$. Lines indicate mean, with shaded areas around them representing 95%-confidence intervals. Big black lines indicate $p \leq 5e^{-2}$. Data and code supporting these findings are available from https://doi.org/10.17605/OSF.IO/SNXQM.

**Supplementary Fig. 5** Correlations between all hypothesis cRSMs used in regressing sensory cRSMs for top-$k$ predictions where $k = 5$. Hypothesis cRSMs included baseline morphs (BM), baseline acoustic predictions (BA), baseline semantic predictions (BS), speaker-invariant acoustic predictions (IA), speaker-specific acoustic predictions (SA), speaker-invariant semantic predictions (IS), and speaker-specific semantic predictions (SS). Baseline predictors were included in all models to control for acoustic properties of the morph (BM) as well as general acoustic (BA) and semantic predictions (BS) that were irrespective of our experiment. Left: Correlations were computed between all hypothesis cRSMs and averaged over time points. Individual cells contain means and standard errors computed over subjects. Right: Correlations were computed between all hypothesis cRSMs, but not averaged over time points. In each cell, time points (0.0-1.0$s$) are plotted on the x-axis, whereas correlations (-1 - 1) are plotted on the y-axis. Per cell, we plot chance level (dashed lines) and mean over subjects (solid lines) with confidence intervals indicated by shaded areas around means. Data and code supporting these findings are available from https://doi.org/10.17605/OSF.IO/SNXQM.

**Supplementary Fig. 6** Validation analysis using foil priors that disrupted participant-specificity of semantic priors. To confirm that performance of top-$k$ acoustic predictions depended on the trial-specific top-$k$ predictions, we compared performance of predictions from the true speaker-invariant and -specific semantic priors with coherent but not trial-specific foil semantic priors in cRSM regressions (see Validating top-$k$ acoustic predictions). In accordance with our main analysis, we set $k = 5$. As expected, this confirmed that speaker-specific acoustic predictions depended on the trial-specific predictions, indicating that they captured meaningful acoustic predictions. Note that, for visual clarity, we restricted the y-axis to a smaller range. Individual bars represent group means, with black lines indicating 95%-confidence intervals around the mean. Big bars between groups indicate $p \leq 5e^{-2}$. Data and code supporting these findings are available from https://doi.org/10.17605/OSF.IO/SNXQM.

**Supplementary Fig. 7** Between-item RSA regression models using $k = 1$ predictions to validate results using a conventional approach and without spectrogram averaging (see Conventional representational similarity analysis). Results confirm the dominance of acoustic sharpening (sh) over acoustic prediction errors (pe), as well as the dominance of acoustic over semantic sharpening. Unlike the within-item cRSM regression, this approach also shows modest additive variance explained by both acoustic prediction errors and semantic sharpening and prediction errors, though variance uniquely attributable to either is comparatively smaller (see Supplementary Table 7). Therefore, this does not change our interpretation that acoustic sharpening is the dominant computation at the sensory level, but suggests that multiple computations may co-occur at the same level to varying degrees. Individual bars represent group means, with black lines indicating 95%-confidence intervals around the mean. ***, **, * $p \leq 1e^{-3}$, $p \leq 1e^{-2}$, and $p \leq 5e^{-2}$ for model comparisons against baseline, respectively. Big bars between models indicate $p \leq 5e^{-2}$. Data and code supporting these findings are available from https://doi.org/10.17605/OSF.IO/SNXQM.

**Supplementary Fig. 8** Between-item RSA regression models, setting $k = 1$ to avoid averaging and restricting the temporal window of the analysis to the shortest possible prediction (0.0-0.48$s$) such that differences in length could not affect results (see Conventional representational similarity analysis). Results corroborate the pattern observed without temporal restrictions (see Supplementary Fig. 7), ruling out prediction length as a confounding factor (see Supplementary Table 8). Individual bars represent group means, with black lines indicating 95%-confidence intervals around the mean. ***, **, * indicate $p \leq 1e^{-3}$, $p \leq 1e^{-2}$, and $p \leq 5e^{-2}$, respectively. Big bars between groups indicate $p \leq 5e^{-2}$. Data and code supporting these findings are available from https://doi.org/10.17605/OSF.IO/SNXQM.

**Supplementary Fig. 9** To identify the layer within word2vec2.0 that best captured phonotactic, lexical and semantic surprisal, we performed back-to-back decoding[2] over an independent set of narrative stimuli (see Pretrained transformers as statistical surrogates). Here, we show the relative causal contribution of each feature across layers, for a number of PCA projections with dimensionality $d$. Small dots represent means, with 95%-confidence intervals around them. Downward-facing triangles represent the best layer overall for each $d$. Crucially, for all $d$ tested, layer 19 (transformer layer 12) was identified as the most informative layer. This is in accordance with recent results finding that middle layers of transformers tend to reflect neural processing best[3]. Data and code supporting these findings are available from https://doi.org/10.17605/OSF.IO/SNXQM.

**Supplementary Fig. 10** To test the robustness of the effect of speaker-specific semantic surprisal, we refit the models using $d = 10$ features from transformer activations. Left: This revealed that, again, only speaker-specific semantic surprisal improved encoding performance—though here results varied more strongly, likely due to the now relatively high number of predictors within the model. Big dots represent group means, with 95%-confidence intervals around them. Small dots represent individual participants. ***, **, * indicate $p \leq 1e^{-3}$, $p \leq 1e^{-2}$, and $p \leq 5e^{-2}$, respectively. The downward-facing triangle marks the best model overall. Big bold lines indicate $p \leq 5e^{-2}$.

Right: As before, speaker-specific semantic surprisal explained significant variance, with a cluster across a wide array of sensors around $110ms$-$400ms$. The inlaid topography shows variance explained at each sensor position within the cluster, with channels contributing to the cluster highlighted in black. Lines indicate means, with 95%-confidence intervals as shaded areas around them. Bold black lines indicate $p \leq 5e^{-2}$. Data and code supporting these findings are available from https://doi.org/10.17605/OSF.IO/SNXQM.

**Supplementary Fig. 11** To further probe the robustness of speaker-specific semantic surprisal, we refit the models using control predictors derived from the word that was more likely under the speaker prior. Consequently, these models were implicitly biassed against speaker-specific semantic surprisal, as this disambiguation of the morph meant some speaker-specific information was already encoded in the control models. Left: Again, we find that speaker-specific semantic surprisal was the only predictor that significantly improved model performance. Big dots represent group means, with 95%-confidence intervals around them. Small dots represent individual participants. ***, **, and * represent $p \leq 1e^{-3}$, $p \leq 1e^{-2}$, and $p \leq 5e^{-2}$, respectively. The downward-facing triangle marks the best model overall. Big black lines indicate $p \leq 5e^{-2}$. Right: Again, we find that speaker-specific semantic surprisal explains significant variance, with a cluster across all sensors between $-100ms$-$410ms$. The inlaid topography shows variance explained at each sensor position, with sensors contributing to the cluster marked in black. Lines indicate means, with 95%-confidence intervals around them. Big bold lines indicate $p \leq 5e^{-2}$. Data and code supporting these findings are available from https://doi.org/10.17605/OSF.IO/SNXQM.

**Supplementary Fig. 12** Peak latencies of semantic surprisal effects across morph, congruent and incongruent trials, demonstrating that there was no significant difference between peak latencies. Note that these represent raw latency estimates over the temporal extent of the cluster. No jackknife procedure was applied. Big dots indicate group means, with 95%-confidence intervals around them. Small dots represent individual participants. Data and code supporting these findings are available from https://doi.org/10.17605/OSF.IO/SNXQM.

**Supplementary Fig. 13** Overview of stimulus materials. **A** Visualisation of the semantic space spanned by the words used in this experiment, as projected into two dimensions using PCA. Dots represent individual words, coloured by the semantic context they were most likely under. **B** Gaussian approximation of the distribution of perceptual bias of morphs across the experiment, showing that stimuli exhibit no systematic bias. **C** Perceptual bias of morphs for individual contexts. Big dots indicate context means, with 95%-confidence intervals around them. Small dots represent individual morphs. Again, no systematic biases exist. Data and code supporting these findings are available from https://doi.org/10.17605/OSF.IO/SNXQM.

**Supplementary Table 1** Results from generalised linear model over responses from online experiment.

**Supplementary Table 2** Results from generalised linear model over responses from EEG experiment.

**Supplementary Table 3** Results from stimulus reconstructions for each frequency band.

**Supplementary Table 4** Results from consecutive contrasts of top-$k$ within-item cRSM regression models.

**Supplementary Table 5** Results from contrasts in within-item cRSM regression models using $k = 19$.

**Supplementary Table 6** Results from within-item cRSM regression models, comparing foil priors to real participant-specific priors (see Validating top-$k$ acoustic predictions).

**Supplementary Table 7** Results from between-item RSA regression models using $k = 1$ (see Conventional representational similarity analysis).

**Supplementary Table 8** Results from between-item RSA regression models using $k = 1$ and restricting the temporal window to 0.0-0.48$s$ to rule out prediction length as a confounding factor (see Conventional representational similarity analysis).

**Supplementary Table 9** Results from contrasts in single-trial encoding models using pretrained transformers with $d = 10$.

**Supplementary Table 10** Results from contrasts in single-trial encoding models using control predictors derived from target words.

**Supplementary Table 11** Results from contrasts in single-trial encoding models in the follow-up task.

**Supplementary Table 12** All stimulus pairs along with their corresponding context pairs used in the present study.
